# Genome-wide CRISPR synthetic lethality screen identifies a role for the ADP-ribosyltransferase PARP14 in replication fork stability controlled by ATR

**DOI:** 10.1101/2020.04.08.032847

**Authors:** Ashna Dhoonmoon, Emily M. Schleicher, Claudia M. Nicolae, Kristen E. Clements, George-Lucian Moldovan

**Affiliations:** Department of Biochemistry and Molecular Biology, The Pennsylvania State University College of Medicine, Hershey, PA 17033, USA

## Abstract

The DNA damage response is essential to maintain genomic stability, suppress replication stress, and protect against carcinogenesis. The ATR-CHK1 pathway is an essential component of this response, which regulates cell cycle progression in the face of replication stress. PARP14 is an ADP-ribosyltransferase with multiple roles in transcription, signaling, and DNA repair. To understand the biological functions of PARP14, we catalogued the genetic components that impact cellular viability upon loss of PARP14 by performing an unbiased, comprehensive, genome-wide CRISPR knockout genetic screen in PARP14-deficient cells. We uncovered the ATR-CHK1 pathway as essential for viability of PARP14-deficient cells, and identified regulation of replication fork stability as an important mechanistic contributor to the synthetic lethality observed. Our work shows that PARP14 is an important modulator of the response to ATR-CHK1 pathway inhibitors.

## Introduction

The DNA damage response (DDR) machinery is essential to maintain genomic stability, ensure cellular proliferation, and protect against carcinogenesis (1). The complex mechanisms employed by the DDR participate not only in repairing DNA damage, but also in attenuating replication stress (2,3). Arrest of the DNA polymerases at sites of replication blockades can result in collapse of the replication machinery and genomic instability. A crucial component of the DDR is the ataxia telangiectasia and Rad3-related (ATR) protein kinase, which is activated by single stranded DNA induced upon replication stress. This leads to downstream phosphorylation of Chk1, which induces a broad cellular response resulting in stabilization of the replication fork, suppression of origin firing, and cell cycle arrest (4-6). ATR and CHK1 inhibitors are currently being investigated as anti-cancer drugs, with multiple clinical trials under way (7,8).

ADP-ribosylation is a prominent post-translational modification which regulates transcription, signal transduction, and DNA repair (9-12). PARP1 is the best characterized member of this family. In particular, PARP1 has emerged as potent target for treatment of tumors with deficiency in the BRCA pathway of homologous recombination (HR) DNA repair, as unrepairable DNA damage results in death of HR-deficient cells treated with PARP1 inhibitors such as olaparib (13,14). This synthetic lethal interaction between PARP1 and the BRCA pathway has been effectively employed for clinical treatment of ovarian cancer (15).

Besides PARP1, the PARP family comprises 16 other members, with various and less understood functions (16). PARP14 (also known as ARTD8) has been associated with multiple cellular processes, however mechanistic details are generally sparse (17). PARP14 has been shown to be involved in regulation of multiple signal transduction pathways including NFκB (18-20), and JNK (21,22). Moreover, PARP14 has been described as a transcriptional co-activator regulating the macrophage-specific transcriptional program (23-25). More recently, it has been shown that PARP14 interacts with multiple RNA regulatory proteins and may play a role in regulating RNA stability (23,26). PARP14 catalytic inhibitors are currently being developed and targeting PARP14 has been proposed as a possible therapeutic approach for multiple cancer types (18,21,22,27,28).

We previously showed that PARP14 is essential for genomic stability by promoting HR and alleviating replication stress (29). Mechanistically, we showed that PARP14 regulates the association of the RAD51 recombinase, an essential HR factor, with damaged DNA. These findings further indicate that PARP14 may impact the tumor response to treatment with genotoxic drugs.

With the advent of the genomics era, and the concomitant development of numerous novel drug targets, it has become clear that identification of the genetic background that confers maximum drug sensitivity is paramount for advancing cancer therapy. Genome-wide genetic screens in human cells have proven invaluable tools to comprehensively and unbiasedly evaluate pharmacogenetic interactions (30-32). Moreover, such screens can provide invaluable insights into functions and mechanisms of human genes. Here, we describe a genome-wide CRISPR-based knockout screen designed to identify synthetic lethality interactions of PARP14. We show that the ATR-CHK1 pathway is essential for viability of PARP14-deficient cells, and identify regulation of replication fork stability as an important mechanistic contributor to the synthetic lethality observed. Our work shows that PARP14 is an important modulator of the response to ATR-CHK1 pathway inhibitors.

## Results

### Genome-wide CRISPR screen identifies PARP14 synthetic lethal candidates

In order to identify genes which are essential for cellular viability in the absence of PARP14, we performed a genome-wide synthetic lethality CRISPR knockout screen in 8988T pancreatic cancer cells (Figure 1A). First, we obtained PARP14-knockout 8988T cells by CRISPR/Cas9-mediated genome editing (Figure 1B). Out of the several PARP14-knockout clones obtained, the KO6 clone (PARP14^KO6^) was used for the synthetic lethality screen. Wildtype and PARP14^KO6^ 8988T cells were infected with the Brunello human CRISPR knockout lentiviral-based library. This library targets 19,114 genes with a total of 76,441 unique guide RNAs (gRNAs), thus on average covering each gene with four different gRNAs (33). To maintain 250-fold library coverage, 20 million library-infected cells were allowed to grow for two weeks. Cells were the collected, and genomic DNA was extracted. The gRNA region was amplified by PCR and identified by Illumina sequencing (Figure 1A).

**Figure 1.**
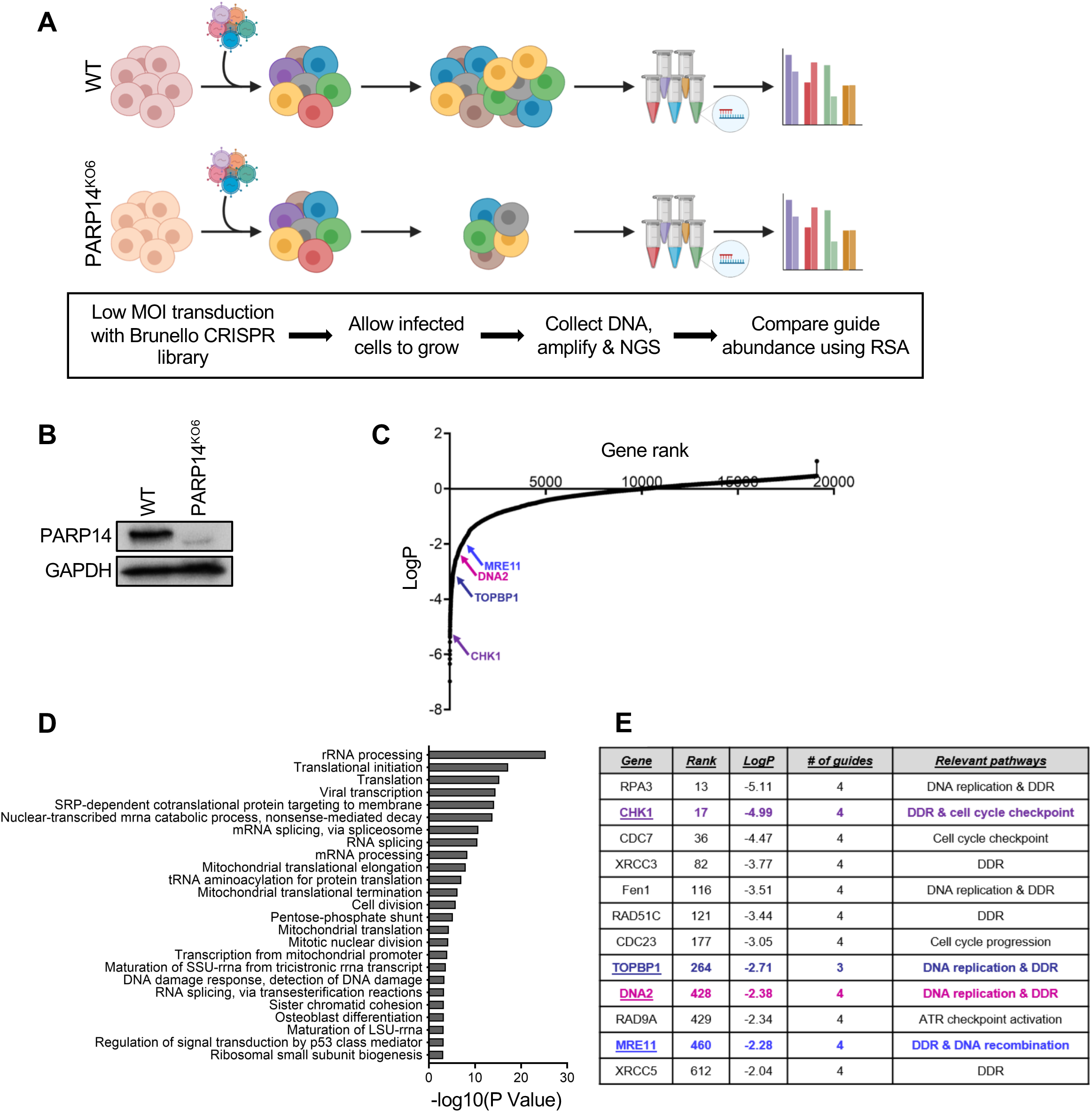
Genome-wide CRISPR knockout screen identified genes essential for viability of PARP14-knockout cells. **(A)** Schematic representation of CRISPR knockout screen. 8988T wild type (WT) and PARP14-knockout (PARP14^KO6^) were infected with the Brunello CRISPR knockout library and allowed to grow for two weeks. Genomic DNA was then extracted from both groups of cells and gRNAs were identified using Illumina sequencing. **(B)** Western blot showing loss of PARP14 protein in the 8988T PARP14^KO6^ cells. **(C)** Scatterplot ranking the genes targeted by library according to P-values is shown. RSA analysis was used to obtain gene ranking. **(D)** Pathway analysis showing the biological process that were significantly enriched in the top 500 hits (genes lost in the PARP14^KO6^ cells compared to wildtype). The top 25 Gene Ontology (GO) terms are shown. **(E)** Multiple DNA damage response (DDR) genes were among the top hits. The highlighted candidates, namely CHK1, DNA2, TOPBP1 and MRE11, were validated in this study.

We next employed the Redundant siRNA Activity (RSA) algorithm (34) to generate a ranked list of genes that were lost in PARP14-knockout compared to wildtype control condition (Figure 1C; Supplemental Table S1). Biological pathway analysis of the top 500 hits revealed RNA-related processes as the most commonly enriched in synthetic lethal interactions with PARP14 loss (Figure 1D, Supplemental Table S2), perhaps in line with previously proposed roles for PARP14 in regulating RNA stability (23,26). Another biological process highly represented on the pathway analysis and previously associated with PARP14 was regulation of mitochondrial activity (21,27). However, cell division, chromosome biology and DNA replication and repair also feature prominently on the list (Figure 1D, E). In particular, among the top hits were multiple components of the ATR pathway, including CHK1, TOPBP1, MRE11, RPA3, and RAD9A (Figure 1E), suggesting an involvement of PARP14 in suppressing replication stress. This is broadly in line with our previous study uncovering a role for PARP14 in DNA repair (29).

### Loss of CHK1 or DNA2 reduces proliferation of PARP14-deficient cells

For screen validation, as a proof of concept we first picked two of the functionally relevant top candidates, namely CHK1 and DNA2. Both CHK1 and DNA2 are key players in DNA damage repair and represent potential therapeutic targets for cancer therapy (35-37). To validate these candidates, we used both the original 8988T PARP14^KO6^ cell line in which the screen was performed, as well as two additional PARP14-knockout 8988T clones, namely PARP14^KO14^ and PARP14^KO19^ (Figure 2A). We employed siRNA to knockdown the candidates in these cells (Figure 2B), and measured their proliferation over four days using the CellTiterGlo ATP-based luminescence assay. Consistent with the screen results, CHK1 knockdown led to impaired cellular proliferation in all three 8988T PARP14-knockout clones compared to wildtype cells (Figure 2C). Similar findings were observed when DNA2 was knocked down (Figure 2D). We also measured apoptosis using Annexin V flow cytometry upon CHK1 depletion in PARP14-knockout cells. Treatment of multiple PARP14-knockout 8988T clones with siRNA targeting CHK1 significantly increased apoptosis compared to control cells (Figure 2E).

**Figure 2.**
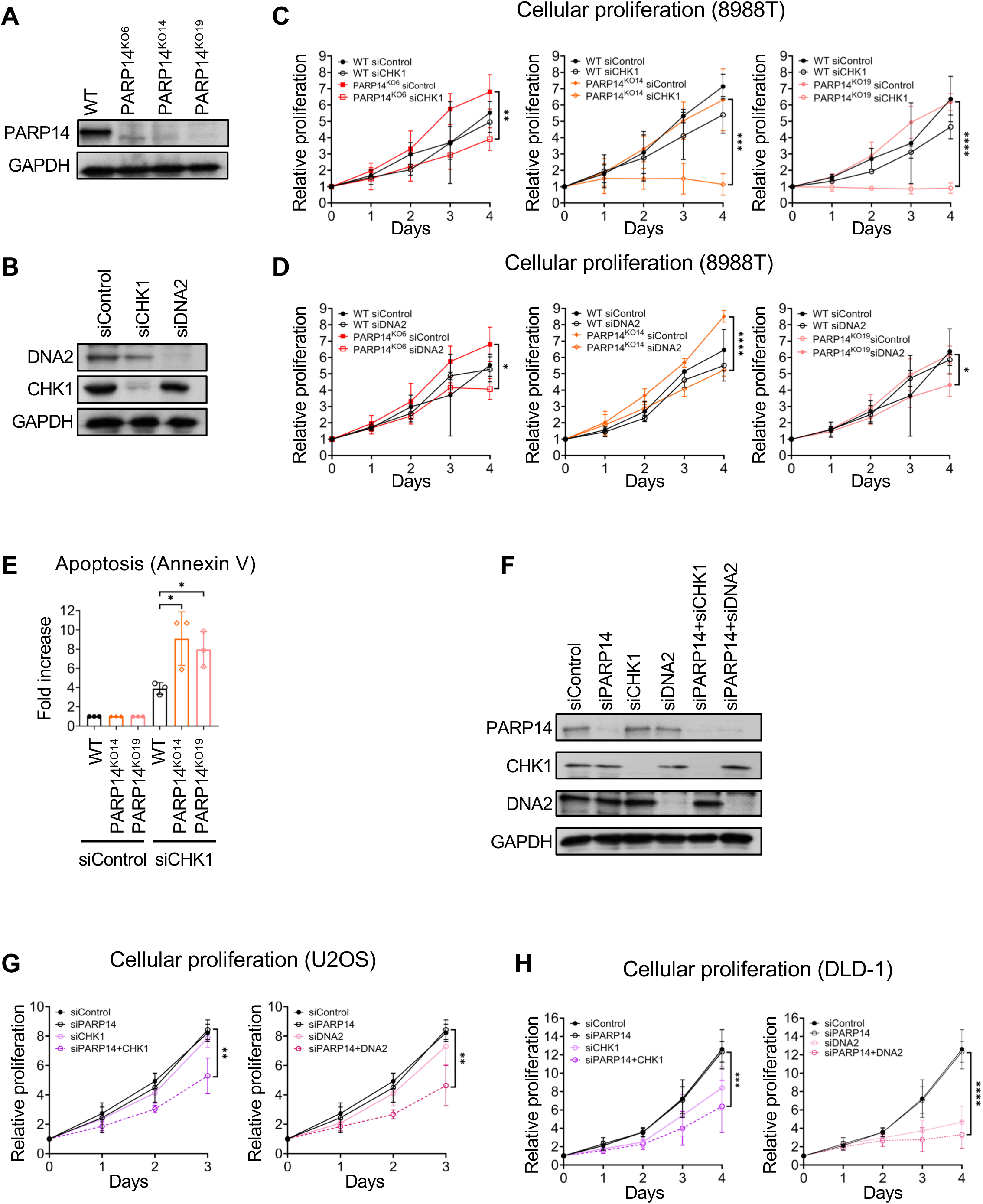
Knockdown of CHK1 or DNA2 reduces proliferation of PARP14-deficient cells. **(A)** Western blot showing the loss of PARP14 protein in multiple 8988T knockout clones. **(B)** Western blot showing efficient siRNA-mediated downregulation of CHK1 and DNA2 in 8988T cells. **(C)** Cellular proliferation assay showing that CHK1 knockdown reduced proliferation of all three PARP14-knockout 8988T clones compared to control. The average of three experiments is presented, with standard deviations shown as error bars. Asterisks indicate statistical significance. **(D)** Cellular proliferation assay showing that DNA2 knockdown reduced proliferation of all three PARP14-knockout 8988T clones compared to control. The average of three experiments is presented, with standard deviations shown as error bars. Asterisks indicate statistical significance. **(E)** Annexin V assays demonstrating increased apoptosis in multiple PARP14-knockout 8988T clones upon CHK1 knockdown. The average of three experiments is presented, with standard deviations shown as error bars. Asterisks indicate statistical significance. **(F)** Western blot showing efficient siRNA-mediated co-depletion of PARP14 and CHK1 or DNA2 in DLD-1 cells. **(G, H)** Co-depletion of CHK1 or DNA2 reduces proliferation of PARP14-knockdown U2OS **(G)** and DLD-1 **(H)** cells. The average of three experiments is presented for U2OS cells and average of four experiments is presented for DLD-1 cells, with standard deviations shown as error bars. Asterisks indicate statistical significance.

In order to rule out any cell line specific effects, we next sought to validate CHK1 and DNA2 in two additional cell lines, namely U2OS (human osteosarcoma) and DLD-1 (colorectal adenocarcinoma). For these two cell lines, we performed co-depletion of PARP14 and either CHK1 or DNA2 using siRNA. Western blot experiments indicated that co-depletion was efficient (Figure 2F). In line with the findings in 8988T cells, loss of both PARP14 and CHK1 or DNA2 reduced cellular proliferation in U2OS (Figure 2G) and DLD-1 (Figure 2H) cell lines. These results indicate that CHK1 and DNA2 are essential for proliferation of PARP14-deficient cells.

Next, we tested how long-term viability is affected when the top candidates are depleted in the PARP14-knockout cells. To this end, we performed clonogenic survival assays in 8988T cells. In all three knockout clones, siRNA-mediated depletion of CHK1 resulted in severely impaired colony formation (Figure 3A, B). To rule out off-target effects of the CRISPR gene editing system employed, we corrected the PARP14^KO6^ clone by exogenous, constitutive re-expression of *PARP14* cDNA. Two separate re-expression clones (#1 and #2) were obtained (Figure 3A). Re-expression of PARP14 in the KO6 clone restored the clonogenic survival upon CHK1 depletion to wildtype levels (Figure 3B). Similar to CHK1, depletion of DNA2 in all three 8988T PARP14-knockout clones also resulted in reduced clonogenic survival, which was rescued upon re-expression of PARP14 cDNA in the PARP14^KO6^ clone (Figure 3C). Moreover, the synthetic lethality interaction between PARP14 and CHK1 or DNA2 was further validated by crystal violet staining of plates seeded at high density with PARP14-knockout cells treated with siRNA targeting these factors (Figure 3D, E). These findings confirm that PARP14 is synthetic lethal with CHK1 and DNA2, thus validating our genome-wide synthetic lethality screen.

**Figure 3.**
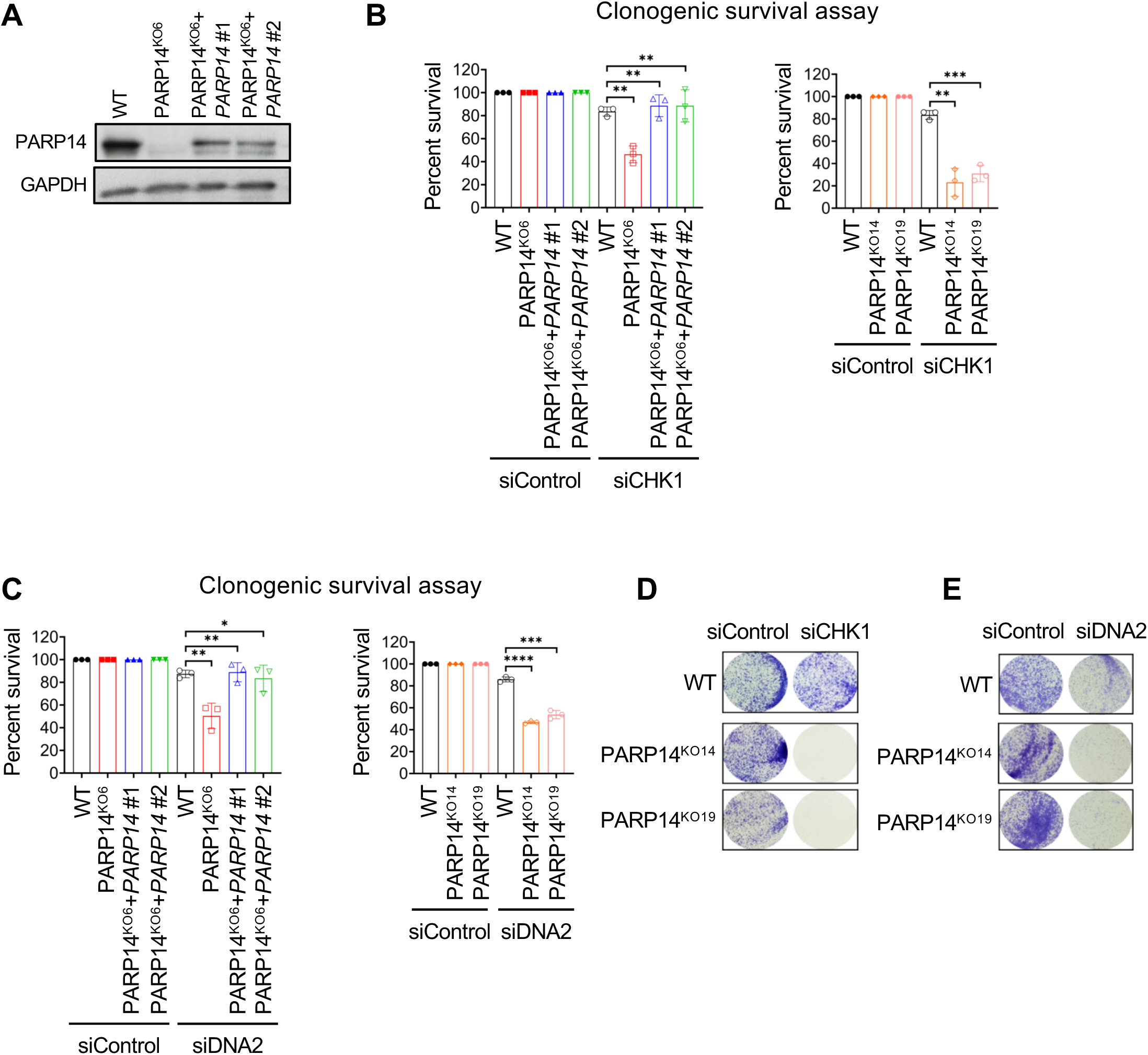
Reduced viability of PARP14-knockout cells upon depletion of CHK1 or DNA2. **(A)** Western blot showing re-expression of PARP14 in the 8988T knockout cells corrected with *PARP14* cDNA. Two different clones were obtained and are investigated here. **(B, C)** Clonogenic survival assays showing reduced survival of PARP14-knockout 8988T cells upon CHK1 **(B)** or DNA2 **(C)** knockdown. All three PARP14-knockout clones were investigated and showed similar phenotypes. Re-expression of exogenous PARP14 in the knockout cells rescued the survival. The average of three experiments is presented, with standard deviations shown as error bars. Asterisks indicate statistical significance. **(D, E)** Representative images of crystal violet staining showing the reduced viability of PARP14-knockout 8988T cells upon depletion of CHK1 **(D)** or DNA2 **(E)**. Two different knockout clones show the same phenotype.

### Synthetic lethality between PARP14 and ATR pathway components

In addition to CHK1, multiple other components of the ATR-CHK1 pathway were among the top hits in our PARP14 synthetic lethality screen, including RPA3, TOPBP1, RAD9A and MRE11 (Figure 1C, E). TOPBP1 and MRE11, which is a member of the MRN complex, co-operate to activate ATR in response to replication stress (38-41). Thus, we decided to also validate these two candidates. Western blot experiments indicated that TOPBP1 can be efficiently depleted from 8988T cells (Figure 4A). Similar to observations made with the other top hits, knocking down TOPBP1 led to impaired colony formation in two different PARP14-knockout 8988T clones (Figure 4B). Moreover, TOPBP1 depletion significantly increased apoptosis in these cells (Figure 4C).

**Figure 4.**
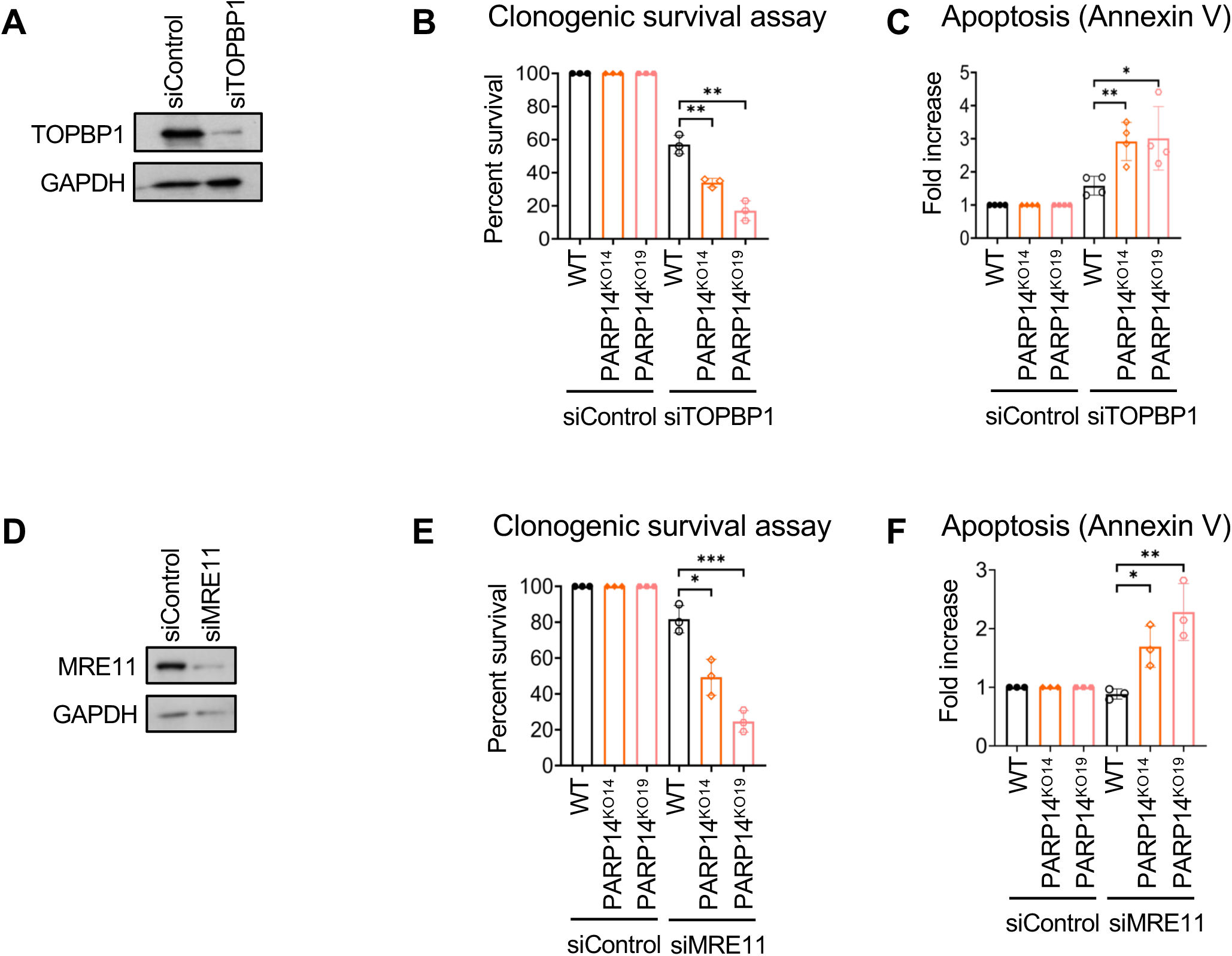
Reduced viability of PARP14-knockout cells upon inactivation of TOPBP1 or MRE11. **(A)** Western blot showing efficient siRNA-mediated downregulation of TOPBP1 in 8988T cells. **(B)** Clonogenic survival assays showing reduced survival of PARP14-knockout 8988T cells upon TOPBP1 knockdown. The average of three experiments is presented, with standard deviations shown as error bars. Asterisks indicate statistical significance. **(C)** Annexin V assays demonstrating increased apoptosis in PARP14-knockout 8988T cells upon TOPBP1 knockdown. The average of four experiments is presented, with standard deviations shown as error bars. Asterisks indicate statistical significance. **(D)** Western blot showing efficient siRNA-mediated downregulation of MRE11 in 8988T cells. **(E)** Clonogenic survival assays showing reduced survival of PARP14-knockout 8988T cells upon MRE11 knockdown. The average of three experiments is presented, with standard deviations shown as error bars. Asterisks indicate statistical significance. **(F)** Annexin V assays demonstrating increased apoptosis in PARP14-knockout 8988T cells upon MRE11 knockdown. The average of three experiments is presented, with standard deviations shown as error bars. Asterisks indicate statistical significance.

Finally, we also depleted MRE11 from 8988T cells (Figure 4D). MRE11 knockdown in two different PARP14-knockout clones resulted in reduced clonogenic survival (Figure 4E), and increased apoptosis (Figure 4F). These findings confirm that TOPBP1 and MRE11, upstream components of the ATR pathway, are required for viability of PARP14-deficient cells. Moreover, these findings further validate our CRISPR knockout screen.

### PARP14-knockout cells show hypersensitivity to pharmacological inhibition of the ATR-CHK1 pathway

Pharmacological inhibition of enzymatic activity is a key approach in personalized cancer therapy. Having observed that CHK1 depletion impairs cellular viability of PARP14-knockout cells, we wanted to confirm these observations using a pharmacological approach. Rabusertib (LY2603618) is a selective CHK1 inhibitor (CHK1i). To test sensitivity of PARP14-deficient cells to CHK1 inhibition, we measured cellular proliferation of PARP14-knockout cells treated with increasing concentrations of rabusertib. Cellular viability of all three PARP14-knockout clones was significantly reduced compared to wildtype control (Figure 5A). Re-expression of exogenous P*ARP14* cDNA in PARP14^KO6^ cell line restored cellular viability (Figure 5A). Similar results were obtained when using clonogenic survival assays (Figure 5B, C).

**Figure 5.**
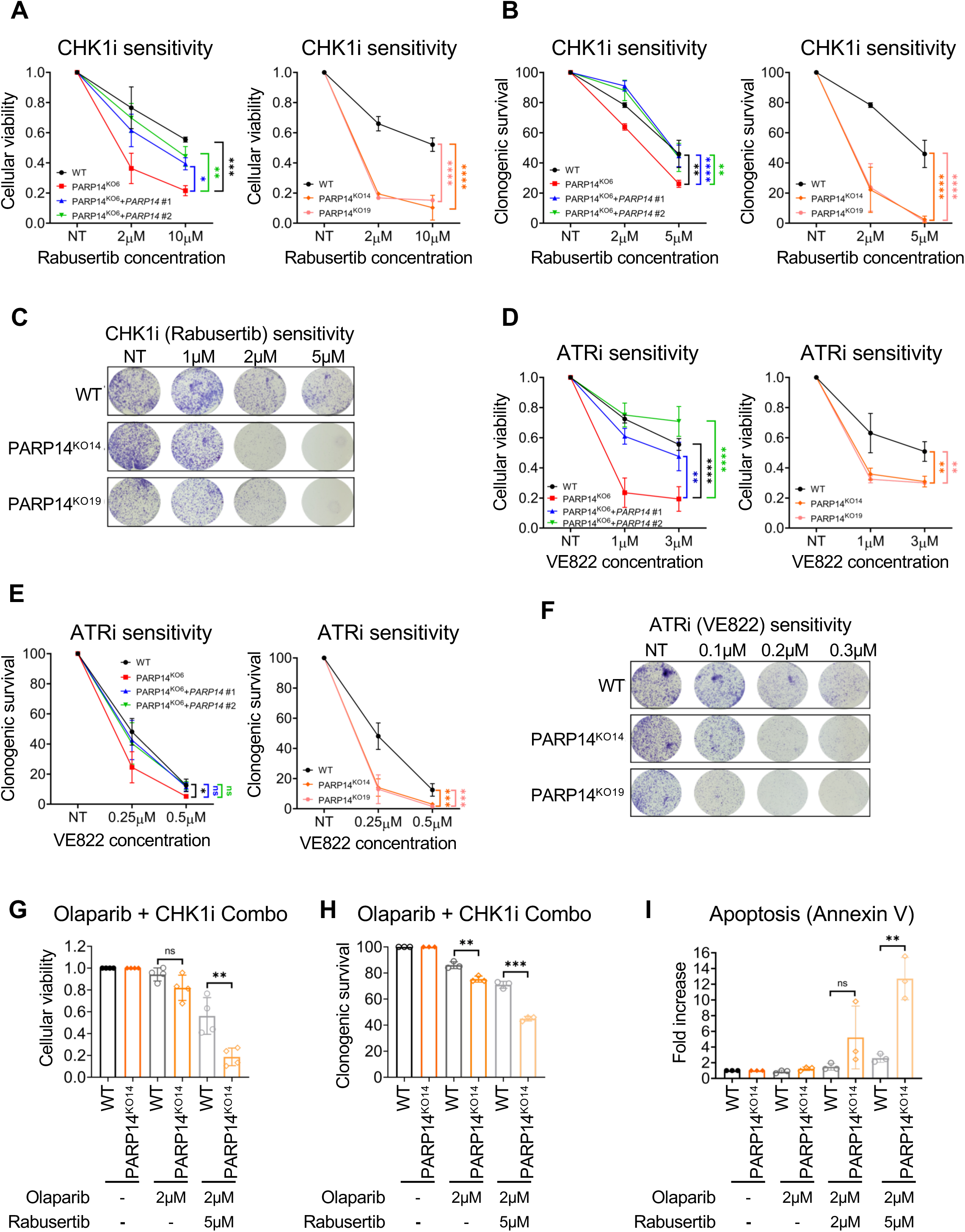
Loss of PARP14 sensitizes cells to inhibitors of the ATR-CHK1 pathway. **(A, B)** Increased sensitivity of 8988T PARP14-knockout cells to the CHK1 inhibitor rabusertib, in both cellular viability **(A)** and clonogenic **(B)** assays. Multiple knockout clones show the same phenotype. Re-expression of PARP14 in the knockout cells restores CHK1i resistance. The average of three experiments is presented, with standard deviations shown as error bars. Asterisks indicate statistical significance. **(C)** Crystal violet staining showing increased rabusertib sensitivity of 8988T PARP14-knockout cells. **(D, E)** Increased sensitivity of 8988T PARP14-knockout cells to the ATR inhibitor VE822, in both cellular viability **(D)** and clonogenic **(E)** assays. Multiple knockout clones show the same phenotype. Re-expression of PARP14 in the knockout cells restores ATRi resistance. The average of three experiments is presented, with standard deviations shown as error bars. Asterisks indicate statistical significance. **(F)** Crystal violet staining showing increased VE822 sensitivity of 8988T PARP14-knockout cells. **(G, H)** CHK1 inhibition potentiates the olaparib sensitivity of PARP14-knockout 8988T cells in both cellular viability **(G)** and clonogenic **(H)** assays. The average of three experiments is presented, with standard deviations shown as error bars. Asterisks indicate statistical significance. **(I)** Annexin V assays demonstrating increased apoptosis in PARP14-knockout 8988T cells upon concomitant treatment with CHK1 and PARP1 inhibitors. The average of three experiments is presented, with standard deviations shown as error bars. Asterisks indicate statistical significance.

Since CHK1 is the key downstream factor in the ATR pathway (4,37), and multiple additional components of this pathway were top candidates in our PARP14 synthetic lethality screen (Figure 1C, E), we sought to investigate whether PARP14-knockout cells are also sensitive to ATR inhibitors (ATRi). To this end, we treated cells with VE822, a selective ATRi. All three PARP14-knockout clones demonstrated higher sensitivity to VE822, in both cellular viability (Figure 5D) and clonogenic survival (Figure 5E, F) assays. This sensitivity was suppressed upon re-expression of wildtype *PARP14* cDNA in the PARP14^KO6^ clone (Figure 5D, E). These findings show that PARP14-deficient cells are sensitive not only to genetic depletion of CHK1, but also to pharmacological inhibition of the ATR-CHK1 pathway.

We previously showed that PARP14 is involved in HR, and thus cells depleted of PARP14 by siRNA show slight sensitivity to the PARP1 inhibitor olaparib (29). We observed a similar trend for the PARP14-knockout 8988T cells in both cellular viability (Figure 5G) and clonogenic (Figure 5H) assays. However, co-treatment with the CHK1 inhibitor rabusertib dramatically increased the olaparib sensitivity of PARP14-knockout cells (Figure 5G, H). Moreover, co-treatment with rabursetib and olaparib significantly increased apoptosis in PARP14-knockout cells compared to control cells (Figure 5I). These findings further attest to the importance of the PARP14 status as an important genetic determinant of the cellular response to cancer drugs targeting the DNA repair system.

### Replication fork stability defects underlie the synthetic lethality between PARP14 and the ATR pathway

Upon replication stress, the ATR-CHK1 pathway promotes replication fork stability, preventing fork collapse and chromosome breakage (42-44). Thus, we investigated replication fork stability by employing the DNA fiber combing assay to measure the progress of individual replication forks, upon consecutive incubations with thymidine analogs IdU and CldU. Immunofluorescence microscopy-based detection of replication tracts indicated that, under normal growth conditions, loss of PARP14 does not affect replication tract length (Figure 6A). However, CHK1 depletion significantly reduced replication tract length in PARP14-knockout cells (Figure 6B). We further validated the knockdown studies by employing pharmaceutical inhibitors of the ATR1-CHK1 pathway. Similar to the knockdown studies, CHK1 inhibition in PARP14-knockout cells resulted in a stronger reduction in replication tract length in PARP14-knockout cells compared to control cells (Figure 6C). Similar results were observed for ATR inhibition (Figure 6D). These results indicate an increased necessity for ATR activation to maintain viability of PARP14-deficient cells, perhaps reflecting increased endogenous replication stress in these cells. In line with this, we observed increased γH2AX in PARP14-knockout cells, both under normal growth conditions and in particular upon ATR inhibition (Figure 6E), indicating that ATR-mediated fork protection suppresses accumulation of abnormal DNA structures in PARP14-deficient cells.

**Figure 6.**
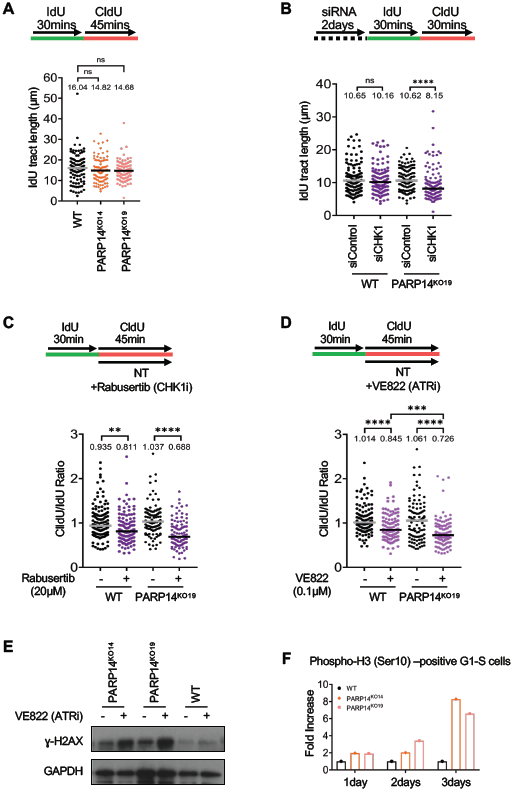
Impact of ATR-CHK1 pathway inhibition on replication dynamics of PARP14-knockout cells. **(A)** DNA fiber combing assay showing normal replication tract length in multiple PARP14-knockout clones. **(B)** Knockdown of CHK1 reduces replication tract length in PARP14-knockout 8988T cells. **(C, D)** Inhibition of CHK1 (**C**) or ATR (**D**) reduces replication tract length in PARP14-knockout 8988T cells. For all DNA fiber experiments (**A-D**), the median values are indicated for each sample, and the asterisks indicate statistical significance. At least 100 fibers were quantified. A schematic representation of the assay is shown at the top. (**E**) Western blot showing that H2AX phosphorylation is increased in PARP14-knockout cells upon treatment with 0.25μM VE822 for 24h. (**F**) Quantification of G1 and S cells with phosphorylated histone H3 at Ser10 as detected by flow cytometry. The fold increase normalized to wildtype is shown upon treatment with 0.25μM VE822 for the indicated number of days (n=1).

The decreased replication fork tracts in PARP14-deficient cells upon ATR inhibition suggest that replication is deficient in these cells, possibly because of fork arrest at endogenous lesions. The ATR pathway is also important for suppressing cell cycle progression in the presence of DNA damage, and its inhibition results in premature mitotic entry of cells with DNA damage (45,46). Histone H3 phosphorylation at Ser10 is a marker of chromosome condensation in mitosis. We observed an increase in G1-S cells positive for H3-Ser10 phosphorylation upon ATR inhibition in PARP14-knockout cells (Figure 6F). Overall, these findings indicate that, upon concomitant loss of PARP14 and the ATR pathway, DNA replication is defective, and cells with incompletely replicated DNA enter mitosis resulting in mitotic catastrophe, thereby explaining the loss of viability.

## Discussion

Genome-wide screens are powerful tools to investigate biological roles of the gene of interest in an unbiased manner. The PARP14 synthetic lethality screen described here identified a number of biological processes with which PARP14 activity has been previously associated. The most highly represented biological processes involved RNA metabolism, in line with previously published work indicating a role for PARP14 in RNA stability (23,26). In addition, this may also reflect a possible activity of PARP14 in directly binding RNA, as PARP14 contains two RRM domains, which are known to bind RNA (16). In addition, biological processes highly enriched among the top hits which were previously associated with PARP14 functions include mitochondrial activity (21,27) and the DDR (29).

Understanding how the specific molecular make-up of the tumor modulates its response to therapy allows improved utilization of cancer drugs. This is relevant for both classic genotoxic chemotherapeutics such as DNA damaging compounds (eg. cisplatin) and replication inhibitors (eg. hydroxyurea), as well as for the new generation of chemical inhibitors such as those targeting the DDR, including PARP1 inhibitors and inhibitors of the ATR-CHK1 pathway. By employing a genome-wide CRISPR knockout screen aimed at unbiased identification of synthetic lethal interactors of PARP14, we found that the ATR-CHK1 pathway was essential for viability of PARP14-deficient cells, in multiple cell lines. PARP14-deficient cells were hypersensitive to both genetic depletion, and pharmacological inhibition of this pathway. Multiple components of the pathway were identified, including the upstream components TOPBP1, MRE11, RPA3 and RAD9A, and the downstream kinase CHK1. Although not a direct component of the ATR pathway, the DNA2 nuclease-helicase, another top hit we validated, has been found to participate in ATR activation under certain conditions, at least in yeast (36,47). These findings highlight an important role of PARP14 in the response to replication stress.

Mechanistically, we identified the control of DNA replication fork stability as a potential contributor to the synthetic lethality between the ATR-CHK1 pathway and PARP14. We show that DNA replication is compromised upon concomitant loss of PARP14 and the ATR pathway, and cells with incompletely replicated DNA undergo premature mitotic entry, thereby explaining the synthetic lethality observed.

The ATR-CHK1 pathway suppresses origin firing, thus its inhibition increases the number of replication forks and decreases the nucleotide pools in the cell, resulting in slower replication fork speed (6,42-44). In PARP14-deficient cells this slowing is accentuated, resulting in severe replication deficiency. We hypothesize that this occurs because of increased stalling of the replication forks at sites of DNA lesions, resulting in accumulation of abnormal DNA structures. However, it is also possible that this reflects a role of PARP14 in replication origin firing. Indeed, another top candidate from the synthetic lethality screen is CDC7 (Figure 1E), a kinase which regulates origin firing (48).

In conclusion, we identified an unexpected role for the ATR-CHK1 pathway in promoting cellular viability in the context of PARP14 deficiency. Our work indicates that the status of the *PARP14* gene in the tumor is an important determinant of the tumor response to DDR inhibitors, which are emerging as a powerful class of cancer drugs.

## Materials and Methods

### Cell culture

Human 8988T and U2OS cells were grown in Dulbecco’s modified Eagle’s medium (DMEM). DLD-1 cells were grown in Roswell Park memorial Institute (RPMI) 1640 medium. DMEM and RPMI were both supplemented with 10% FBS and penicillin/streptomycin. To generate the 8988T PARP14-knockout cells, the commercially available PARP14 CRISPR/Cas9 KO plasmid was used (Santa Cruz Biotechnology sc-402812). Transfected cells were FACS-sorted into 96-well plates using a BD FACSAria II instrument. Resulting colonies were screened by Western blot. To re-express exogenous PARP14 in the knockout cell lines, cells were infected with the lentiviral construct pLV-Puro-SV40>Flag/hPARP14 (Cyagen) was used, constitutively expressing Flag-tagged PARP14 under the control of the SV40 promoter.

Gene knockdown was performed using Lipofectamine RNAiMAX transfection reagent. AllStars Negative Control siRNA (Qiagen 1027281) was used as control. The following oligonucleotide sequences (Stealth siRNA, ThermoFisher) were used:

PARP14: AGGCCGACTGTGACCAGATAGTGAA

DNA2: TTAGAATGCAGGCAACTGTATCCTT

MRE11: CATTACATACCTGCCTCGAGTTATT

TOPBP1: Silencer Select ID s2183

CHK1: Silencer Select ID s504

Denatured whole cell extracts were prepared by boiling cells in 100 mM Tris, 4% SDS, 0.5M β-mercaptoethanol. Antibodies used for Western blot were: PAPR14 (Santa Cruz Biotechnology sc-377150); Chk1 (Cell signaling Technology 2360); DNA2 (Abcam ab96488); MRE11 (Santa Cruz Biotechnology sc-135992); TOPBP1 (Novus NB100-217); GAPDH (Santa Cruz Biotechnology sc-47724); γH2AX (Abcam ab-2893).

The chemical inhibitors used in this study were obtained from Selleck Chemicals: rabusertib (CHK1i); VE822 (ATRi); olaparib (PARP1i).

### CRISPR screens

For CRISPR knockout screens, the Brunello Human CRISPR knockout pooled lentiviral library (Addgene 73179) was used (33). This library targets 19,114 genes with 76,411 gRNAs. 100 million 8988T (wildype and PARP14^KO6^) cells were infected with this library at a multiplicity of infection (MOI) of 0.4 to achieve 500x coverage and selected for 4 days with 1.25 µg/mL puromycin. For each condition, 20 million cells freshly infected with the library (to maintain 250x coverage) were seeded and allowed to grow for two weeks. Genomic DNA was isolated using the DNeasy Blood and Tissue Kit (Qiagen 69504) per the manufacturer’s instructions. gRNAs were amplified using PCR primers with Illumina adapters. Genomic DNA from 20 million cells (250-fold library coverage) was used as template for PCR. The PCR reaction contained 10µg of gDNA, with 20µl 5X HiFi Reaction Buffer, 4µl of P5 primer, 4µl of P7 primer, 3µl of Radiant HiFi Ultra Polymerase (Stellar Scientific), and water. The P5 and P7 primers used were determined using the user guide provided with the CRISPR libraries (https://media.addgene.org/cms/filer_public/61/16/611619f4-0926-4a07-b5c7-e286a8ecf7f5/broadgpp-sequencing-protocol.pdf). The purified PCR product was sequenced with Illumina HiSeq 2500 single read for 50 cycles.

For bioinformatic analysis of the screen results, the custom python script provided (count_spacers.py) (49) was used to calculate sgRNA representation. The difference between the number of guides present in the PARP14-knockout condition compared to the wildtype condition was determined. Specifically, one read count was added to each sgRNA, and then the reads from the PARP14-knockout condition were normalized to the wildtype condition. The values obtained were then used as input in the Redundant siRNA Activity (RSA) algorithm (50). For RSA, the Bonferroni option was used and guides that were at least 2-fold enriched in the PARP14-knockout condition compared to the wildtype condition were considered hits. The p-values are determined by the RSA algorithm for the genes that are most enriched in the PARP14-knockout condition compared to the wildtype condition. Analyses of the Gene Ontology pathways enriched among the top hits was performed using DAVID (51,52).

### Functional cellular assays

For clonogenic survival assays, 500 cells were seeded per well in 6-well plates and treated with siRNA or drug as indicated. Media was changed after 3 days and cells were allowed to grow for 10-14 days. Colonies formed were then washed with PBS, fixed with a solution of 10% methanol + 10% acetic acid and stained with crystal violet (2%, Aqua solutions).

For crystal violet imaging, 50,000 cells were seeded per well in 12-well plates and treated with the appropriate drug concentrations. Staining was performed 3 days later as described above.

To assess cellular proliferation, a luminescent ATP-based assay was performed using the CellTiterGlo reagent (Promega G7572) as per manufacturer’s instructions. Following treatment with siRNA, 1500 cells were seeded per well (day 0) and plates were read daily for 5 days. For drug sensitivity, 1500 cells were seeded per well in 96-well plated and treated with the indicated drug doses. Plates were read 3 days later.

For apoptosis assays, cells were treated with siRNA for 2 days, followed by media change. Cells were prepared for flow cytometry two days after media change using the FITC Annexin V kit (Biolegend, 640906). Quantification was performed using a BD FACSCanto 10 flow cytometer.

For quantification of G1-S cells positive for histone H3 phosphorylated at Ser10, the Click-iT Plus EdU Alexa Fluor 488 Flow Cytometry Assay Kit (ThermoFisher) was used to measure cell cycle distribution, according to the manufacturer’s instruction. Concomitantly, cells were stained with the Phospho-Histone H3 (Ser10) Alexa Fluor 594 conjugated antibody. Cells were subsequently analyzed by flow cytometry.

### DNA fiber assays

For the experiments with gene knockdown, cells were treated with siRNA for 2 days, then incubated with 100µM IdU for 30 min, washed with PBS and incubated with 100 µM CldU. For the experiments with drug treatment, cells were incubated with 100µM IdU for 30mins, washed with PBS, and incubated within the drugs and/or CldU as indicated. Next, cells were collected and processed using the the FiberPrep kit (Genomic Vision EXT-001) according to the manufacturer’s instructions. DNA molecules were stretched onto coverslips (Genomic Vision COV-002-RUO) using the FiberComb Molecular Combing instrument (Genomic Vision MCS-001). Slides were stained with antibodies detecting CldU (Abcam 6236), IdU (BD 347580), and DNA (Millipore Sigma MAD3034). Slides were then incubated with secondary Cy3, Cy5, or BV480-conjugated antibodies (Abcam 6946, Abcam 6565, and BD Biosciences 564879). Finally, the cells were mounted onto coverslips and imaged using a confocal microscope (Leica SP5).

### Statistical analyses

For CellTiter-Glo cellular proliferation assays, the 2-way ANOVA statistical test was used. This test was also used for drug sensitivity clonogenic assay. For clonogenic survival assays upon gene knockdown by siRNA, as well as for the Annexin V assay, the t-test (two-tailed, unequal variance unless indicated) was used. For the DNA fiber assay, the Mann-Whitney statistical test was performed. Statistical significance is indicated for each graph (ns = not significant, for P>0.05; * for P≤0.05; ** for P≤0.01; *** for P≤0.001, **** for P≤0.0001).

## Acknowledgements

We would like to thank Drs. Hong-Gang Wang and Robert Brosh for materials and advice; and the following Penn State College of Medicine core facilities: Flow Cytometry, Genomic Analyses, and Imaging. This work was supported by: NIH R01ES026184 and R01GM134681 (to GLM) and NIH 1F31CA243301 (to EMS).

**Legends to Supplemental Tables**

**Supplemental Table S1**. Lists of all genes in the CRISPR knockout screen ranked by p-value.

**Supplemental Table S2**. List of the top 25 Gene Ontology terms from the pathway analysis of the top 500 hits.

## Notes

### Competing Interest Statement

The authors have declared no competing interest.

## References

1. Ciccia, A. and Elledge, S.J. (2010) The DNA damage response: making it safe to play with knives. Mol Cell, 40, 179–204.

2. Zeman, M.K. and Cimprich, K.A. (2014) Causes and consequences of replication stress. Nat Cell Biol, 16, 2–9.

3. Gaillard, H., Garcia-Muse, T. and Aguilera, A. (2015) Replication stress and cancer. Nat Rev Cancer, 15, 276–289.

4. Brown, E.J. and Baltimore, D. (2003) Essential and dispensable roles of ATR in cell cycle arrest and genome maintenance. Genes Dev, 17, 615–628.

5. Marechal, A. and Zou, L. (2013) DNA damage sensing by the ATM and ATR kinases. Cold Spring Harbor perspectives in biology, 5.

6. Saldivar, J.C., Cortez, D. and Cimprich, K.A. (2017) The essential kinase ATR: ensuring faithful duplication of a challenging genome. Nat Rev Mol Cell Biol, 18, 622–636.

7. Fokas, E., Prevo, R., Hammond, E.M., Brunner, T.B., McKenna, W.G. and Muschel, R.J. (2014) Targeting ATR in DNA damage response and cancer therapeutics. Cancer Treat Rev, 40, 109–117.

8. Toledo, L.I., Murga, M. and Fernandez-Capetillo, O. (2011) Targeting ATR and Chk1 kinases for cancer treatment: a new model for new (and old) drugs. Mol Oncol, 5, 368–373.

9. Gibson, B.A. and Kraus, W.L. (2012) New insights into the molecular and cellular functions of poly(ADP-ribose) and PARPs. Nat Rev Mol Cell Biol, 13, 411–424.

10. Feijs, K.L., Forst, A.H., Verheugd, P. and Luscher, B. (2013) Macrodomain-containing proteins: regulating new intracellular functions of mono(ADP-ribosyl)ation. Nat Rev Mol Cell Biol, 14, 443–451.

11. Feijs, K.L., Verheugd, P. and Luscher, B. (2013) Expanding functions of intracellular resident mono-ADP-ribosylation in cell physiology. FEBS J, 280, 3519–3529.

12. Kalisch, T., Ame, J.C., Dantzer, F. and Schreiber, V. (2012) New readers and interpretations of poly(ADP-ribosyl)ation. Trends Biochem Sci, 37, 381–390.

13. Bryant, H.E., Schultz, N., Thomas, H.D., Parker, K.M., Flower, D., Lopez, E., Kyle, S., Meuth, M., Curtin, N.J. and Helleday, T. (2005) Specific killing of BRCA2-deficient tumours with inhibitors of poly(ADP-ribose) polymerase. Nature, 434, 913–917.

14. Farmer, H., McCabe, N., Lord, C.J., Tutt, A.N., Johnson, D.A., Richardson, T.B., Santarosa, M., Dillon, K.J., Hickson, I., Knights, C. et al. (2005) Targeting the DNA repair defect in BRCA mutant cells as a therapeutic strategy. Nature, 434, 917–921.

15. Moore, K., Colombo, N., Scambia, G., Kim, B.G., Oaknin, A., Friedlander, M., Lisyanskaya, A., Floquet, A., Leary, A., Sonke, G.S. et al. (2018) Maintenance Olaparib in Patients with Newly Diagnosed Advanced Ovarian Cancer. N Engl J Med.

16. Otto, H., Reche, P.A., Bazan, F., Dittmar, K., Haag, F. and Koch-Nolte, F. (2005) In silico characterization of the family of PARP-like poly(ADP-ribosyl)transferases (pARTs). BMC Genomics, 6, 139.

17. Schweiker, S.S., Tauber, A.L., Sherry, M.E. and Levonis, S.M. (2018) Structure, Function and Inhibition of Poly(ADP-ribose)polymerase, Member 14 (PARP14). Mini Rev Med Chem, 18, 1659–1669.

18. Yao, N., Chen, Q., Shi, W., Tang, L. and Fu, Y. (2019) PARP14 promotes the proliferation and gemcitabine chemoresistance of pancreatic cancer cells through activation of NF-kappaB pathway. Mol Carcinog, 58, 1291–1302.

19. Feijs, K.L., Kleine, H., Braczynski, A., Forst, A.H., Herzog, N., Verheugd, P., Linzen, U., Kremmer, E. and Luscher, B. (2013) ARTD10 substrate identification on protein microarrays: regulation of GSK3beta by mono-ADP-ribosylation. Cell Commun Signal, 11, 5.

20. Verheugd, P., Forst, A.H., Milke, L., Herzog, N., Feijs, K.L., Kremmer, E., Kleine, H. and Luscher, B. (2013) Regulation of NF-kappaB signalling by the mono-ADP-ribosyltransferase ARTD10. Nat Commun, 4, 1683.

21. Iansante, V., Choy, P.M., Fung, S.W., Liu, Y., Chai, J.G., Dyson, J., Del Rio, A., D’Santos, C., Williams, R., Chokshi, S. et al. (2015) PARP14 promotes the Warburg effect in hepatocellular carcinoma by inhibiting JNK1-dependent PKM2 phosphorylation and activation. Nat Commun, 6, 7882.

22. Barbarulo, A., Iansante, V., Chaidos, A., Naresh, K., Rahemtulla, A., Franzoso, G., Karadimitris, A., Haskard, D.O., Papa, S. and Bubici, C. (2013) Poly(ADP-ribose) polymerase family member 14 (PARP14) is a novel effector of the JNK2-dependent pro-survival signal in multiple myeloma. Oncogene, 32, 4231–4242.

23. Iqbal, M.B., Johns, M., Cao, J., Liu, Y., Yu, S.C., Hyde, G.D., Laffan, M.A., Marchese, F.P., Cho, S.H., Clark, A.R. et al. (2014) PARP-14 combines with tristetraprolin in the selective posttranscriptional control of macrophage tissue factor expression. Blood, 124, 3646–3655.

24. Iwata, H., Goettsch, C., Sharma, A., Ricchiuto, P., Goh, W.W., Halu, A., Yamada, I., Yoshida, H., Hara, T., Wei, M. et al. (2016) PARP9 and PARP14 cross-regulate macrophage activation via STAT1 ADP-ribosylation. Nat Commun, 7, 12849.

25. Mehrotra, P., Riley, J.P., Patel, R., Li, F., Voss, L. and Goenka, S. (2011) PARP-14 functions as a transcriptional switch for Stat6-dependent gene activation. J Biol Chem, 286, 1767–1776.

26. Carter-O’Connell, I., Vermehren-Schmaedick, A., Jin, H., Morgan, R.K., David, L.L. and Cohen, M.S. (2018) Combining Chemical Genetics with Proximity-Dependent Labeling Reveals Cellular Targets of Poly(ADP-ribose) Polymerase 14 (PARP14). ACS chemical biology, 13, 2841–2848.

27. Cho, S.H., Ahn, A.K., Bhargava, P., Lee, C.H., Eischen, C.M., McGuinness, O. and Boothby, M. (2011) Glycolytic rate and lymphomagenesis depend on PARP14, an ADP ribosyltransferase of the B aggressive lymphoma (BAL) family. Proc Natl Acad Sci U S A, 108, 15972–15977.

28. Holechek, J., Lease, R., Thorsell, A.G., Karlberg, T., McCadden, C., Grant, R., Keen, A., Callahan, E., Schuler, H. and Ferraris, D. (2018) Design, synthesis and evaluation of potent and selective inhibitors of mono-(ADP-ribosyl)transferases PARP10 and PARP14. Bioorg Med Chem Lett, 28, 2050–2054.

29. Nicolae, C.M., Aho, E.R., Choe, K.N., Constantin, D., Hu, H.J., Lee, D., Myung, K. and Moldovan, G.L. (2015) A novel role for the mono-ADP-ribosyltransferase PARP14/ARTD8 in promoting homologous recombination and protecting against replication stress. Nucleic Acids Res, 43, 3143–3153.

30. Zimmermann, M., Murina, O., Reijns, M.A.M., Agathanggelou, A., Challis, R., Tarnauskaite, Z., Muir, M., Fluteau, A., Aregger, M., McEwan, A. et al. (2018) CRISPR screens identify genomic ribonucleotides as a source of PARP-trapping lesions. Nature, 559, 285–289.

31. Hustedt, N., Alvarez-Quilon, A., McEwan, A., Yuan, J.Y., Cho, T., Koob, L., Hart, T. and Durocher, D. (2019) A consensus set of genetic vulnerabilities to ATR inhibition. Open Biol, 9, 190156.

32. He, Y.J., Meghani, K., Caron, M.C., Yang, C., Ronato, D.A., Bian, J., Sharma, A., Moore, J., Niraj, J., Detappe, A. et al. (2018) DYNLL1 binds to MRE11 to limit DNA end resection in BRCA1-deficient cells. Nature, 563, 522–526.

33. Doench, J.G., Fusi, N., Sullender, M., Hegde, M., Vaimberg, E.W., Donovan, K.F., Smith, I., Tothova, Z., Wilen, C., Orchard, R. et al. (2016) Optimized sgRNA design to maximize activity and minimize off-target effects of CRISPR-Cas9. Nat Biotechnol, 34, 184–191.

34. Konig, R., Chiang, C.Y., Tu, B.P., Yan, S.F., DeJesus, P.D., Romero, A., Bergauer, T., Orth, A., Krueger, U., Zhou, Y. et al. (2007) A probability-based approach for the analysis of large-scale RNAi screens. Nat Methods, 4, 847–849.

35. Jia, P.P., Junaid, M., Ma, Y.B., Ahmad, F., Jia, Y.F., Li, W.G. and Pei, D.S. (2017) Role of human DNA2 (hDNA2) as a potential target for cancer and other diseases: A systematic review. DNA repair, 59, 9–19.

36. Pawlowska, E., Szczepanska, J. and Blasiak, J. (2017) DNA2-An Important Player in DNA Damage Response or Just Another DNA Maintenance Protein? Int J Mol Sci, 18.

37. Rundle, S., Bradbury, A., Drew, Y. and Curtin, N.J. (2017) Targeting the ATR-CHK1 Axis in Cancer Therapy. Cancers (Basel), 9.

38. Burrows, A.E. and Elledge, S.J. (2008) How ATR turns on: TopBP1 goes on ATRIP with ATR. Genes Dev, 22, 1416–1421.

39. Duursma, A.M., Driscoll, R., Elias, J.E. and Cimprich, K.A. (2013) A role for the MRN complex in ATR activation via TOPBP1 recruitment. Mol Cell, 50, 116–122.

40. Olson, E., Nievera, C.J., Lee, A.Y., Chen, L. and Wu, X. (2007) The Mre11-Rad50-Nbs1 complex acts both upstream and downstream of ataxia telangiectasia mutated and Rad3-related protein (ATR) to regulate the S-phase checkpoint following UV treatment. J Biol Chem, 282, 22939–22952.

41. Regal, J.A., Festerling, T.A., Buis, J.M. and Ferguson, D.O. (2013) Disease-associated MRE11 mutants impact ATM/ATR DNA damage signaling by distinct mechanisms. Hum Mol Genet, 22, 5146–5159.

42. Zhong, Y., Nellimoottil, T., Peace, J.M., Knott, S.R., Villwock, S.K., Yee, J.M., Jancuska, J.M., Rege, S., Tecklenburg, M., Sclafani, R.A. et al. (2013) The level of origin firing inversely affects the rate of replication fork progression. The Journal of cell biology, 201, 373–383.

43. Moiseeva, T.N., Yin, Y., Calderon, M.J., Qian, C., Schamus-Haynes, S., Sugitani, N., Osmanbeyoglu, H.U., Rothenberg, E., Watkins, S.C. and Bakkenist, C.J. (2019) An ATR and CHK1 kinase signaling mechanism that limits origin firing during unperturbed DNA replication. Proc Natl Acad Sci U S A, 116, 13374–13383.

44. Mutreja, K., Krietsch, J., Hess, J., Ursich, S., Berti, M., Roessler, F.K., Zellweger, R., Patra, M., Gasser, G. and Lopes, M. (2018) ATR-Mediated Global Fork Slowing and Reversal Assist Fork Traverse and Prevent Chromosomal Breakage at DNA Interstrand Cross-Links. Cell Rep, 24, 2629–2642 e2625.

45. Brown, E.J. and Baltimore, D. (2000) ATR disruption leads to chromosomal fragmentation and early embryonic lethality. Genes Dev, 14, 397–402.

46. Nghiem, P., Park, P.K., Kim, Y., Vaziri, C. and Schreiber, S.L. (2001) ATR inhibition selectively sensitizes G1 checkpoint-deficient cells to lethal premature chromatin condensation. Proc Natl Acad Sci U S A, 98, 9092–9097.

47. Zheng, L., Meng, Y., Campbell, J.L. and Shen, B. (2020) Multiple roles of DNA2 nuclease/helicase in DNA metabolism, genome stability and human diseases. Nucleic Acids Res, 48, 16–35.

48. Moiseeva, T., Hood, B., Schamus, S., O’Connor, M.J., Conrads, T.P. and Bakkenist, C.J. (2017) ATR kinase inhibition induces unscheduled origin firing through a Cdc7-dependent association between GINS and And-1. Nat Commun, 8, 1392.

49. Joung, J., Konermann, S., Gootenberg, J.S., Abudayyeh, O.O., Platt, R.J., Brigham, M.D., Sanjana, N.E. and Zhang, F. (2017) Genome-scale CRISPR-Cas9 knockout and transcriptional activation screening. Nature protocols, 12, 828–863.

50. Birmingham, A., Selfors, L.M., Forster, T., Wrobel, D., Kennedy, C.J., Shanks, E., Santoyo-Lopez, J., Dunican, D.J., Long, A., Kelleher, D. et al. (2009) Statistical methods for analysis of high-throughput RNA interference screens. Nat Methods, 6, 569–575.

51. Ashburner, M., Ball, C.A., Blake, J.A., Botstein, D., Butler, H., Cherry, J.M., Davis, A.P., Dolinski, K., Dwight, S.S., Eppig, J.T. et al. (2000) Gene ontology: tool for the unification of biology. The Gene Ontology Consortium. Nat Genet, 25, 25–29.

52. Huang da, W., Sherman, B.T. and Lempicki, R.A. (2009) Systematic and integrative analysis of large gene lists using DAVID bioinformatics resources. Nature protocols, 4, 44–57.

